# Effect of acute low oxygen exposure on the proliferation rate, viability and gene expression of C_2_C_12_ myoblasts *in vitro*

**DOI:** 10.1101/2020.07.09.162123

**Authors:** Jack V Sharkey

## Abstract

**INTRODUCTION:** Changes in the oxygen concentration of cellular microenvironments play a significant role in regulating cell function during muscle regeneration. Generally, most *in-vitro* cell culture experiments have been carried out in atmospheric conditions with 21% O_2_, which compared to the actual micro-environment of mature skeletal muscle of between 1% and 10% pO2 is extremely hyperoxic (Li et al, 2007). Culturing skeletal muscle cells *in vitro* within their typical physiologically hypoxic environment *in situ* (2-10% pO2) has been shown to increase proliferation rate, reduce apoptosis and increase multiple MRF gene expressions, compared to culturing in a normoxic environment (21% O_2_). However, chronic exposure (>24 hr) to a semi-severe hypoxic environment (≤5% O_2_) can lead to a decrease in cell proliferation and differentiation (Chakravarthy et al, 2001). The effects of acute hypoxic exposure (24 h) has limited research and could be important in understanding the effects of hypoxia on skeletal muscle during brief exposures such as those observed within intermittent hypoxic training programmes. The purpose of this work was to examine the role of acute hypoxia (24 h) on C_2_C_12_ proliferation and relevant gene expression in 2D culture.

**METHODS:** C_2_C_12_ mouse myoblast cells were seeded into six well plates. The cells were maintained in DMEM with 20% FCS. C_2_C_12_ myoblasts were either exposed to 21% or 5% O_2_. The effects of acute hypoxic exposure (24hours) at different time points during the proliferative phase of myogenesis, rather than chronic exposure, on cell proliferation, cellular viability and myogenic regulatory factor gene expression was examined. At 24, 48, 72 and 96 hours RNA was extracted using the Trizol^®^ method and mRNA expression of myogenic regulatory factors, myoD, myf5 and myogenin were detected using the 2-ΔΔCT method. Cell counts and cell viability were also quantified

**RESULTS:** No significant difference was found between cells cultured in normoxic conditions (21%) and those that were exposed to low oxygen conditions for 24hours at various time points over a 96 hour culture period, with regards to proliferation rate, cell viability, and myogenic regulatory gene expression (*Myf5, MyoD* and *Myogenin*).

**DISCUSSION:** The effect of acute low oxygen exposure lasting 24hours appears to not be insufficient in having an effect on the proliferation rate, viability or transcription factor expression of C2C12 cells during the proliferative phase of myogenesis.

## INTRODUCTION

Within human tissue, oxygen concentrations are considerably less than the 21% O_2_ concentrations found in ambient air, and is termed physiological hypoxia. Typically the partial pressure of oxygen (pO_2_) of human tissue lies between 1% and 14% (Stamati, Mudera and Cheema, 2011), with considerable local and regional variations (Figure.1). Within the tissues’ own microenvironment, changes in oxygen concentrations play a significant role in regulating cell function, and can have effects including maintenance of stem cell pluripotency, angiogenesis, stem cell differentiation, ATP regulation, and pH regulation. (Stamati, Tamati and Mudera, 2011; Weidemann *et al*, 2008; Lin *et al*, 2008).

**Figure 1.**
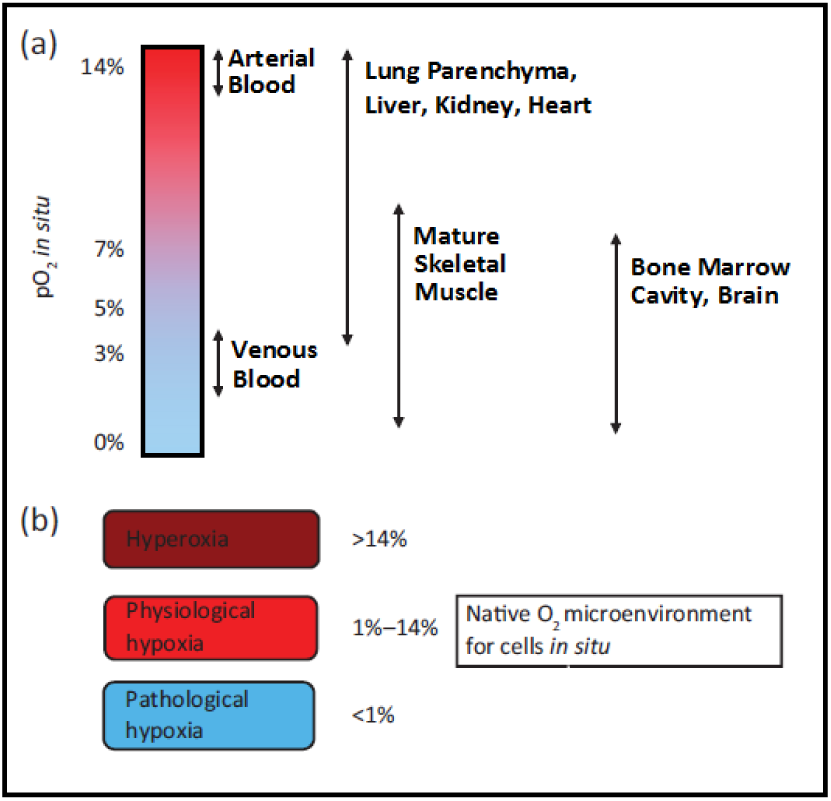
(a) Values of micro-environmental O_2_ found in tissues *in situ* (b) O_2_ micro-environment for cells residing within different tissues in mammalian body. •Image adapted from Stamati, Mudera & Cheema (2011); values taken from Warkerntin (2002), Das, Jahr & Van Osch (2010) & Li, Zhu, Chen & Fan (2007).

Satellite cells, are muscle-resident stem cells found below the basal lamina adhered to the muscle fibres (Mauro, 1961). Activation of the quiescent satellite cells is brought about in order to regenerate or repair muscle tissue, which is generally due to a stress induced change in their micro-environment, caused by either weight bearing exercise or injury (Kuang & Rudnicki, 2008; Miller, Schaefer & Dominov, 1999; Bischoff and Heintz, 1994). Once activated the stem cells undergo proliferation at which point they are known as adult myoblasts. These then differentiate and fuse into multinucleated skeletal muscle fibres (Le Grand & Rudnicki, 2007, Kuang *et al*, 2008). This process, known as myogenesis, is dependent on several transcription factors known as myogenic regulatory factors (MRFs); *MyoD, Myf5, Myogenin* and *MRF4* (Majmundar *et al*, 2012; Cornelsion *et al*, 2000; Charge & Rudnicki, 2004, Yablonka-Reuveni and Rivera, 1994). Prior to activation the quiescent satellite cells express the transcription factor *PAX*_*7*_ (Le Grand & Rudnicki, 2007). Once activated within 12hours there is a rapid up-regulation of *MyoD* and *Myf5*, along with *PAX*_*7*_ and all possess distinct functional roles (Majmndar et al, 2012). *Myf5* is a regulator for muscle progenitor proliferation, whilst *MyoD* is necessary for the consequent differentiation of these precursors (Majmundar et al, 2012). During the proliferation phase of myogenesis, there is co-expression of both *MyoD* and *Myf5* (Cornelison & Wold, 1997).

With the co-existence of *MyoD* and *Myf5*, the genes activate the upregulation of *Myogenin. MyoD* and *Myogenin* stimulate the myoblasts to undergo terminal differentiation and fusion into myofibers through the activation of genes in mature muscle such as the Myosin heavy chain (Kuang et al, 2008; Yablonka-Reuveni and Rivera 1994). Subsequently *P21*, a cell cycle arrest protein, is activated, causing an irreversible exit of the myoblast from the cell cycle. It is also known that *MRF4* is actively expressed during the differentiation phase and continues to be released once newly regenerated myofibers have been fused, indicating that *MRF4* has a role in myofiber maturation. (Zhou and Bornemann, 2001) (Figure.2).

**Figure 2.**
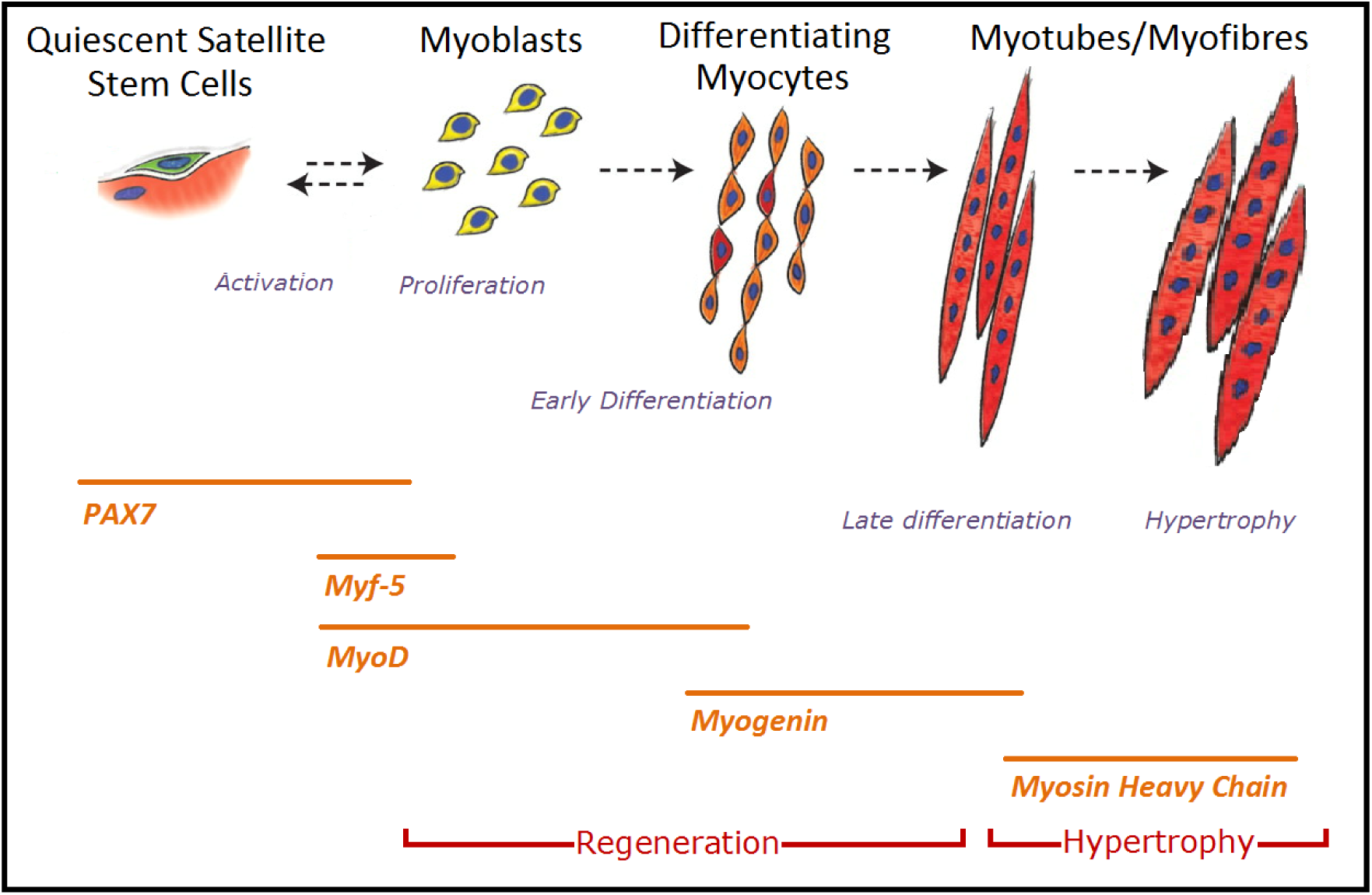
(a) Schematic of the stages of myogenesis from quiescent satellite cells to myotube formaton and hypertrophy, along with an indication of the time periods at which various genes are expressed •Image adapted from Bentzinger, Wang & Rudnicki (2012)

The local microenvironment, otherwise known as the stem cell niche, within the skeletal muscle that accommodates quiescent satellite cells is generally between 1% and 10% pO_2_ (Li 2007). Stress induced changes of the oxygen concentration caused by trauma or weight bearing exercise is believed to be one of the main physiological regulators of satellite cell activity and activation, indicating manipulation of pO2 exposure could be an affective means of improving muscle regeneration (Lui *et al*, 2012). There is some physiological data of local interstitial pO_2_ which has been evaluated using a number of techniques, including oxygen microelectrodes, microspectrophotometry, proton madnetic resonance spectroscopy and phosphorescence quenching (Pittman, 2011., Poole et al, 2011). However, the exact systemic environment of satellite cells that precisely regulates quiescence or activation, and self-renewal or differentiation, is largely unknown (Lui, 2012).

Generally, most *in-vitro* cell culture experiments have been carried out in atmospheric conditions with 21% O_2_, which compared to the actual micro-environment of mature skeletal muscle of between 1% and 10% pO2 is extremely hyperoxic (Li et al, 2007). To develop a better understanding of the molecular mechanisms behind myogenesis and the cell cycle *in-vitro*, it has been suggested that experiments should be performed in environmentally low conditions that mimic the normal physiological environments. Research in this area has shown that satellite cells cultured in physiologically hypoxic conditions (2-10% O_2_) result in a significant increase in proliferation rate, improved survival of mature fibers, decreased satellite cell adipogenisis, promoted myogenesis and larger myotube formations, compared to cultures grown in 21% pO2 (Zhao *et al*, 2003; Chakravarthy *et al*, 2001). It is also noted that the signalling pathways responsible for regulating stem cell proliferation and differentiation are impacted by the O2 concentration of the surrounding micro-environment. With regards to myogenesis and MRF expression, during the proliferation phase there is a significant up-regulation of *MyoD, Myf5* and *Myogenin* when C2C12 murine cells are cultured in 6% O2 conditions compared to 20%. (Csete et al 2001).

When the pO_2_ is dropped below 2% within the microenvironment, also known as pathological hypoxia, the expression of differentiation markers (*MyoD, Myf5*, and *Myogenin*) begin to be inhibited, along with the inhibition of quiescient satellite cell activation and of multinucleated myotube formation (Roy *et al*, 2003). Furthermore there is a notable reduction in satellite cell proliferation and increased apoptosis (Sato *et al*, 2011; Di Carlo *et al*, 2004). The inhibition of myogenic differentiation is found to be through accelerated *MyoD* degredation by the ubiquitin-proteasome pathway (DiCarlo *et al*, 2004).

In general the atmospheric conditions used in traditional muscle cells culture *in vitro* is largely ignored, and recent experiments have found culturing cells in more physiologically atmospheric conditions may be more beneficial for myogenesis and lead to a better understanding of the molecular processes behind skeletal muscle repair and regeneration (Reecy *et al*, 2003). Whether acute exposure of low oxygen on muscular regeneration is as beneficial as chronic exposure is yet to be examined, and may play a vital role in the use of using the manipulation of O_2_ concentrations in improving muscle self renewal, damage repair and treating regenerative disease (Li *et al*, 2007). The aim of the following study is to further support the ever-growing research that culturing cells in a physiologically hypoxic environment is optimal for myogenesis compared to an ambient atmosphere. Specifically the following study aims to establish the effects of twenty-four hours of acute physiological hypoxia (5% pO_2_) during the proliferative phase of myogenesis, including proliferation rate, cellular viability and myogenic regulatory factor gene expression of proliferating C2C12 myoblasts.

## METHOD

### Cell Culture

C2C12 murine myoblasts were provided by the Health Protection Agency Culture Collections (HPA Cultures, Salisbury, UK). Three separate lines of C2C12’s within 1.8ml vial’s were removed from liquid nitrogen and quickly defrosted. The cells were plated at 12500 cells/cm^2^ in T80 culture flasks (80cm^2^ Flasks, NUNC, Denmark) and suspended in 20ml of growth medium (GM) consisting of high glucose Dulbecco’s Modified Eagle’s Medium (DMEM) (Sigma-Aldrich, Haverhill, UK), 20% Foetal Bovine Serum (FBS) (PAA, Paisley, UK), and 1% penicillin/streptomycin (Sigma-Aldrich, Haverhill, UK). Once suspended the cells were kept within a humidified incubator (Heracell 240i CO_2_ Incubator, Thermo Fisher Scientific Inc, USA) where conditions were kept constant at 37°C in a normoxic environment (21% O_2_, 5% CO_2_, 74% N_2_). Growth medium was replaced every 48 hours, and once cells had reached 80% confluence, cells were passaged.

To ensure the line of C2C12 cells used within this experiment could undergo differentiation, a T80 flask was seeded at a density of 12500 cells/cm^2^ and cultured for 96 hours until 100% confluent. Once confluency had been achieved the GM was replaced with differentiation media (DM) to promote myoblast fusion (2% FCS, 1% P/S, and DMEM) (Sigma-Aldrich, Haverhill, UK) and left to culture for a further 48 hours. The cells were then fixed and stained for immunocytochemistry (Figure.3)

**Figure. 3.**
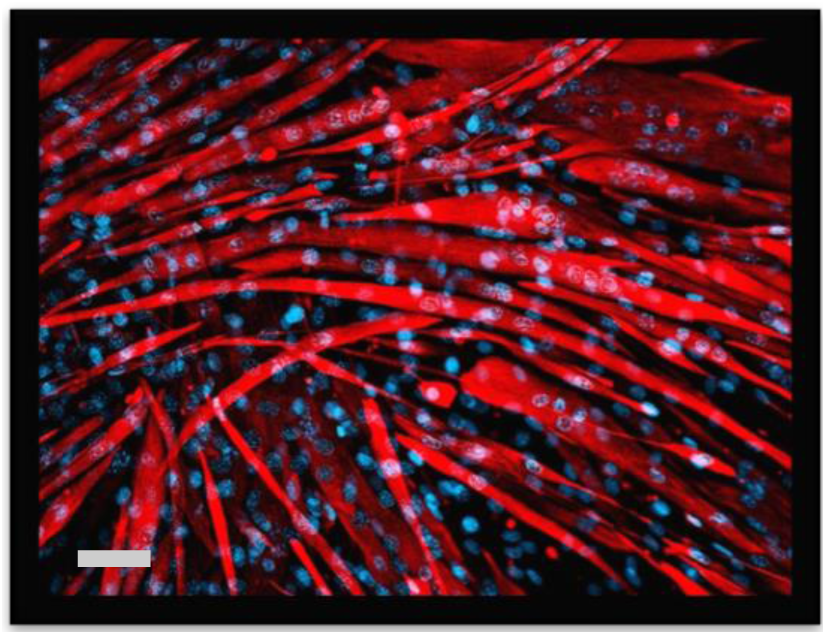
Murine C2C12’s in culture immunostained for desmin (red) with a DAPI nuclear counter stain (blue). Note the formation of myotubes indicating the line of C2C12’s used were able to undergo fusion into multinucleated myotubes, Scale bar = 20µm.

Once a passage number of seven was attained the cells were split and seeded into eight, six well plates (NUNC A/S, Denmark) in 2ml of Growth medium at a density of 1500 cells/cm^2^. The plate layout consisted of three wells for qPCR, one well for immunocytochemistry containing three 13mm sterile glass coverslips, one well for cell counts and microscopy, and one well for checking the oxygen concentration within the microenvironment of the C2C12’s. (Figure.4)

**Figure. 4.**
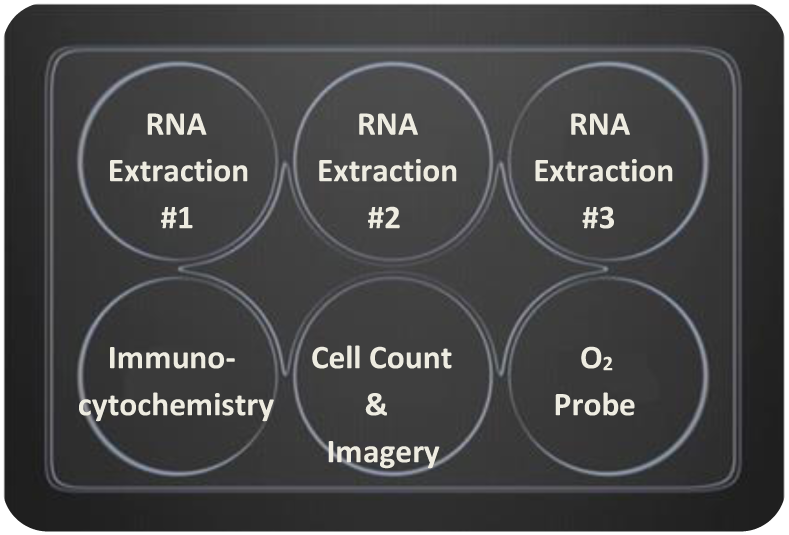
Representation of the six well plate layout.

Cells were sampled for RNA and immunostaining every 24 hours upto a 96 hour timepoint. During this time the C2C12 cells were cultured in a humidified incubator (Heracell 240i CO_2_ Incubator, Thermo Fisher Scientific Inc, USA) at 37°C under normoxic conditions (21% pO_2_). The hypoxic treatment involved an acute exposure of hypoxia during the final twenty-four hours of culture under what would be expected to be a physiological hypoxic condition (5% pO_2_). This was achieved by placing the cells within a SANYO CO_2_/O_2_ hypoxic incubator (MCO-5M, Panasonic, USA) at 37°C (5% O_2_, 5% CO_2_, 90% N_2_). Once the plates were removed from their incubators, they were sampled immediately. A schematic of the experimental design is shown in Figure.5. The GM was refreshed within the plates every twenty-four hours, prior to being placed in either the normoxic or hypoxic incubators. The experiment was repeated three times using three separate lines of C2C12’s.

**Figure. 5.**
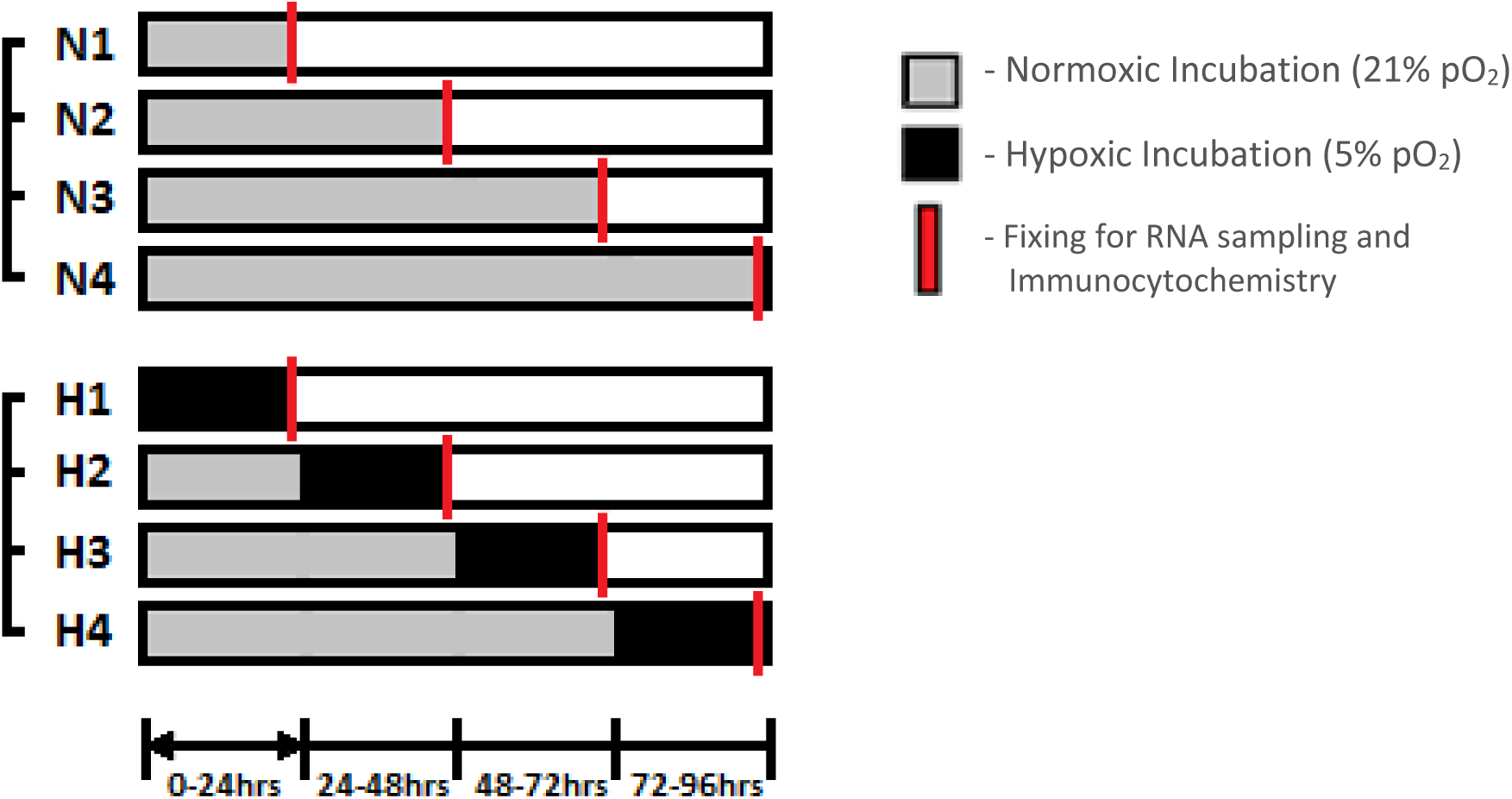
Schematic diagram of the study design N = Normoxic Exposure H = Normoxic with 24hr hypoxic exposure.

To ensure the microenvironment in which the cells were cultured was at a concentration of 5% O_2_ during the hypoxic exposure, a dip-type oxygen microelectrode (MI-730 O_2_ Microelectrode, Microelectrodes Inc, Bedford, UK) along with PowerLab 4/25T with LabChart Software (ADinstruments, New Zealand) was used. The probe was calibrated according to manufacture instructions and placed within the GM of one of the C2C12 inhabited wells through a drilled hole on the roof of the plate. Oxygen concentrations were checked at the end of each twenty-four hour time period for 3 minutes to ensure a constant 5% concentration of oxygen had been reached.

### Cell Counts and Microscopy

The hypoxic and normoxic exposed cells were removed from their incubators and analysed using an LED microscope (Leica DM IL LED, Leica Microsystems, Wetzler, Germany) at twenty-four hour time points throughout the four day culture period and ten images were taken within the designated well for imagery. A cell count of viable cells was performed using a heamoctyometer with Trypan blue exclusion. This was achieved by removing the GM from the monolayer of cells and adding 1ml of Trypsin. Warm Phosphate Buffered Saline solution (PBS) was not placed on the cells so a count of the non-viable adhered cells could be carried out. Once the Trypsin was added, the plate was incubated at 37°C for five minutes with occasional disturbance to displace the cells from the wells. Using the LED microscope the cells were checked for detachment followed by adding 2ml of growth media (GM) to neutralise the effect of the Trypsin. If cells were still confluent, the serum was carefully triturated using a pipette or passed through a filter.

### Immunocytochemistry

Within one of the wells of the six well plates, C2C12 cells were cultured on three 13mm sterile glass coverslips (Sigma-Aldrich, Haverhill, UK). At the end of the designated exposure time in either the normoxic or hypoxic chamber, cells were fixed using ice-cold fixative solution consisting of one part methanol, to one part acetone (1ml). The monolayer of cells were permeabilized using a solution of 1x TRIS buffered saline solution (TBS) along with 5% normal goat serum (GS) and 0.2% Triton X-100 for 30 minutes. Following a thorough wash with TBS, a rabbit monoclonal anti-mouse desmin primary antibody (Sigma-Aldrich, Haverhill, UK) diluted 1 in 200 in TBS solution with 2% GS and 0.2% Triton X-100 was added to the cells and left at room temperature for two hours. A thorough TBS wash ensued before being treated with a goat anti-rabbit TRITC secondary antibody (Sigma-Aldrich, Haverhill, UK) diluted 1 in 200 in TBS with 2% GS and 0.2% Triton X-100 and left for one hour in room temperature with no light exposure. Finally, following a TBS wash, a fluorescent minor-groove DNA binding probe DAPI (4,6-diamidino-2-phenylindole; 1.0ng/ml; Sigma Aldrich, Haverhill, UK) was added to act as a nuclear stain and left for 10 minutes. Once completed the coverslips were thoroughly washed using TBS, dried, before being mounted on glass microscope slides using MOWIOL mounting medium. Examination was carried out by immunoflouresent microscopy using a Leica DM2500 microscope and Leica software LAC V3.7 (Leica Microsystems, Wetzler, Germany).

### RNA Extraction and Quantitative PCR

Levels of proliferation and the onset of differentiation of the 2D culture was determined by quantitative PCR (qPCR) analysis of *MyoD, Myf5*, and *Myogenin* expression. Total RNA was extracted from the cells using the TRIzol^®^ reagent protocol according to manufacturer instructions (Invitrogen). qPCR was run using the Stratagene Mx3005P PCR System (Applied Biosystems, Warrington UK). The PCR reaction mixture (10µl SYBR Green, 0.2µl RT mix, 0.15µl forward primer, 0.15µl reverse primer) was created for each gene to be analysed in duplicate over a 96 well plate (Applied Biosystems). Of this reaction mix,10.5µl of solution was added to each well along with 9.5µl of solution containing 70ng of RNA dissolved in RNAse-free water. The plate was incubated at 50°C for 10 minutes and 95°C for 5 minutes before undergoing 40 cycles of 95°C for 10 seconds and 60°C for 30 seconds, a thermal profile specifically designed by Applied Biosystems for use with assays designed according to Applied Biosystems assay design guidelines. The primer sequences used are presented in Table.1.

**Table 1.**
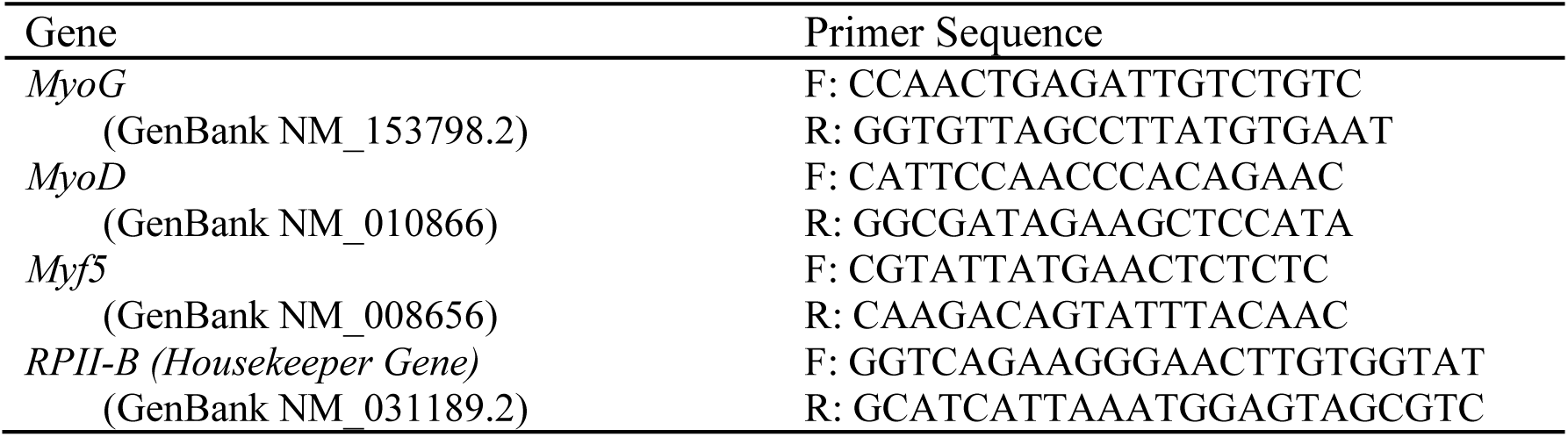
Primer sequences used for PCR analysis. MyoG - Myogenin Factor 4, MyoD – Myogenic Differentiation 1, Myf5 – Myogenic Factor 5.

A cycle threshold (C_T_) was defined as the fractional cycle number at which the fluorescence generated by cleavage of the probe exceeded a fixed threshold above the baseline. A Comparative C_T_ method outlined by Applied Biosystems, was used to quantitate the amount of each target gene present. The mean C_T_ values of the duplicate samples from each sample was determined and normalised to RPII, which acted as the endogenous housekeeping gene. Relative quantitation of mRNA expression was calculated using the ddCT method, using the 2^-ΔΔCt^formula (Livak & Schmittgen, 2001).

### Statistical Analysis

A mixed measures ANOVA was performed to assess differences across treatments (Hypoxia and Normoxia) and across time (24h, 48h, 72h, and 96h). Paired *t*-tests were used to compare between Normoxia and Hypoxia treatment differences at individual time points. Significance was accepted when *P*<0.05. Statistical tests were conducted using SPSS version 19.0 (SPSS, Inc, Chicago, Illinois, USA). Data are reported as mean ± SD.

## RESULTS

### Cell Counts and Viability

The Heamocytometer assay with Trypan blue exclusion showed no significant difference in proliferation rates between cells cultured under normoxia or acute hypoxia at any of the twenty four hour time points (Figure.6). However, collectively over time, there was a notable increase in Total Cell Count with significant differences between time points. There was no significant difference in cell counts between twenty-four hours and forty-eight hours, although cell count increased significantly over time between 48 and 72 hours (P<0.026), and between 48 and 96 hours (P<0.003). There was a notable rise in cell counts between seventy-two hours and ninety-six hours of culturing but this was found to be non-significant (P<0.126).

**Figure. 6.**
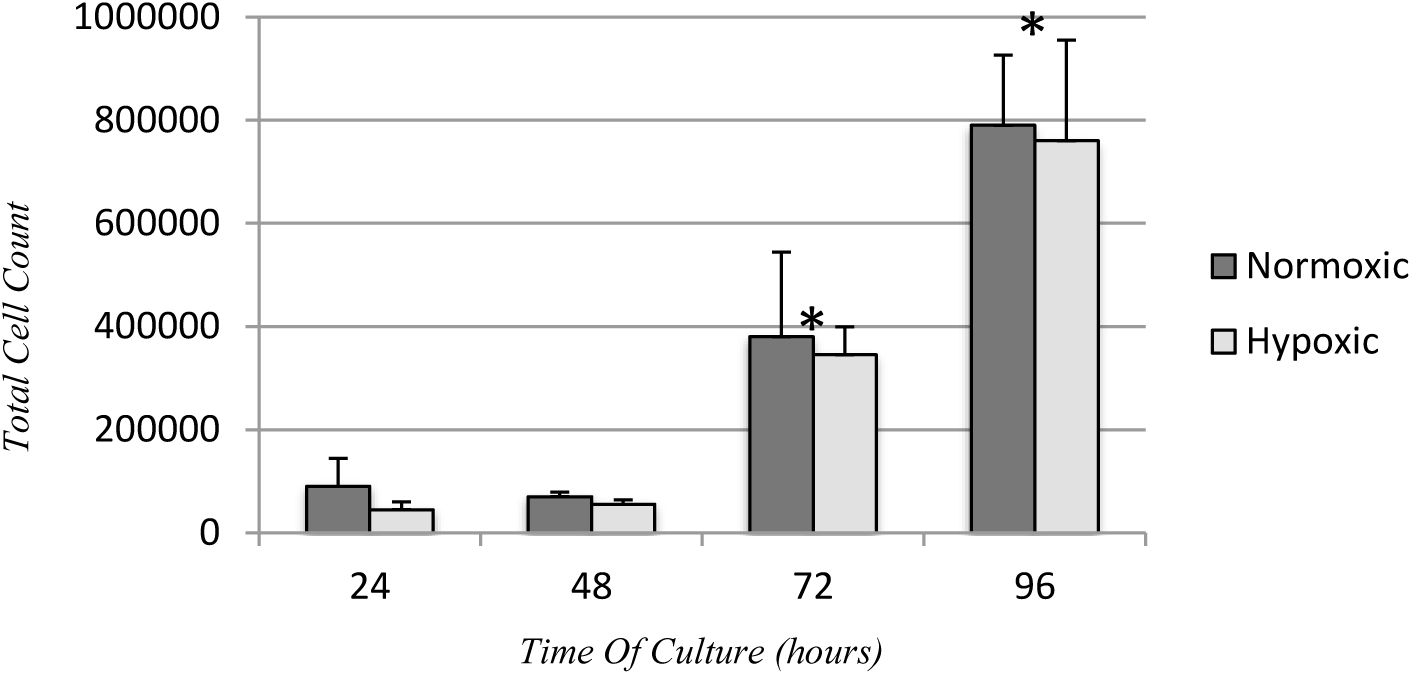
Total cell count of adhered cells, taken every 24 hours over a 96 hour period in vitro, as determined by a haemocytometer. Values expressed as mean ± standard deviation. No significant differences were observed between Normoxic and Hypoxic exposure at any time point with total cell counts (P >0.05).

The percentage of viable cells within the total cell count did not differ significantly at any of the twenty four hour time points between cells cultured under normoxia or acute hypoxia. Although there is a possible trend towards a reduction in cell viability after forty-eight hours before recovering at seventy-two and ninety-six hours. Over time, cell viability did not differ between any of the twenty-four hour time points over the ninety-six hour culture period (Figure.7)

**Figure. 7.**
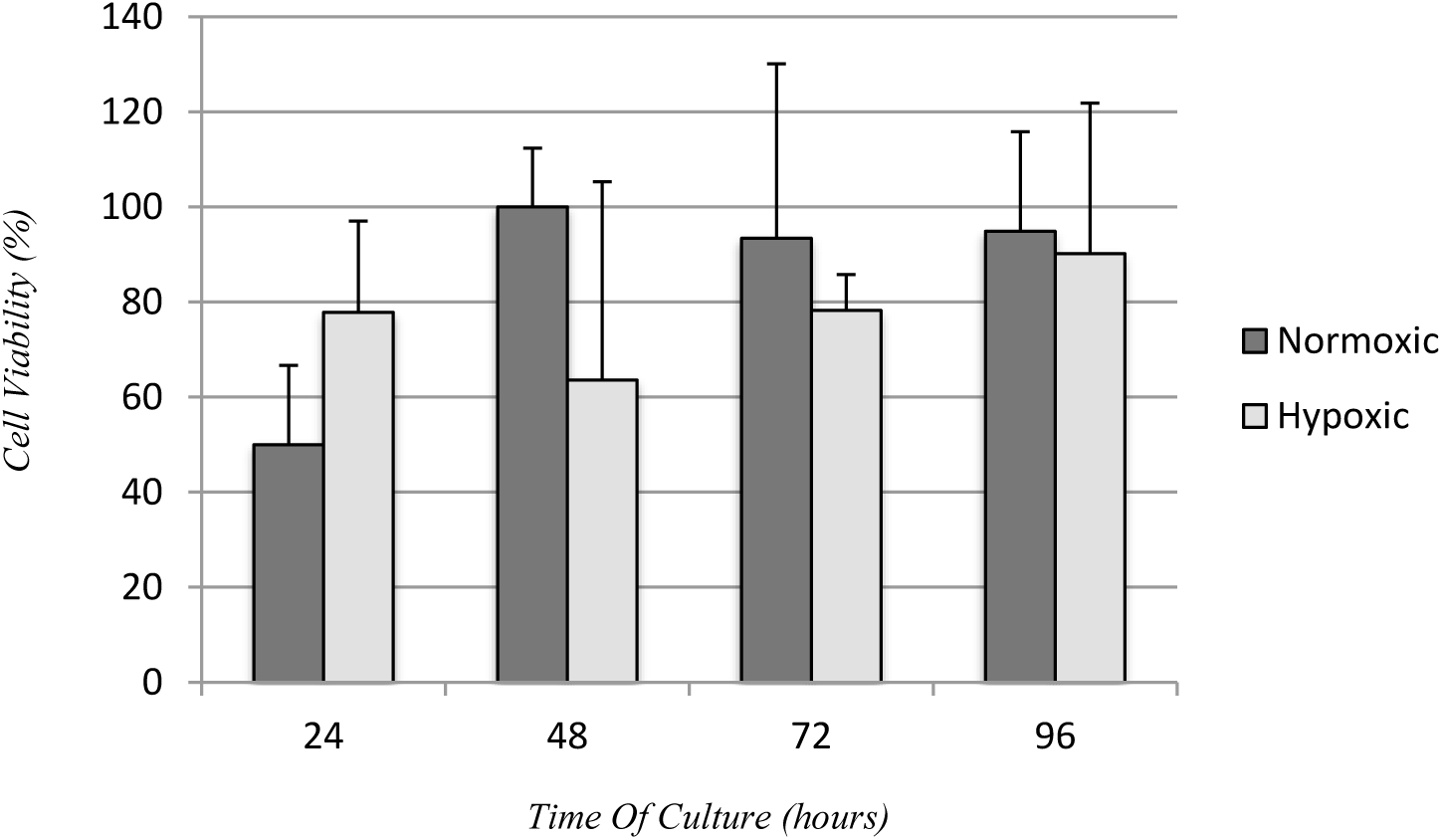
Percentage of viable cells adhered to coverslips taken every 24 hours over a 96 hour period in vitro, as determined by haemocytometer and Trypan blue exclusion. Hypoxic treatment involved acute hypoxic exposure of 5% O_2_ during the final 24 hours of culturing. Values expressed as mean ± standard deviation..

### Quantitative PCR Analysis

Quantative PCR analysis found no significant difference in relative *Myf5, MyoD* and *Myogenin* mRNA expression levels between the two exposure conditions of normoxia or acute hypoxia following twenty-four, forty-eight, seventy-two and ninety-six hours of culturing (P>0,05) (Figure.8). Concerning changes in mRNA expression levels over time, no differences were found between any of the time points when analysing *Myf5* and *MyoD*, however there does appear to be a trend toward *Myf5* expression increasing over time in both conditions. *Myogenin* mRNA expression levels following twenty-four hours of culturing were significantly higher than those expressed following forty-eight hours (P<0.06), seventy-two (P<0.011) and ninety-six (P<0.014) hours of culturing with both treatments.

**Figure. 8.**
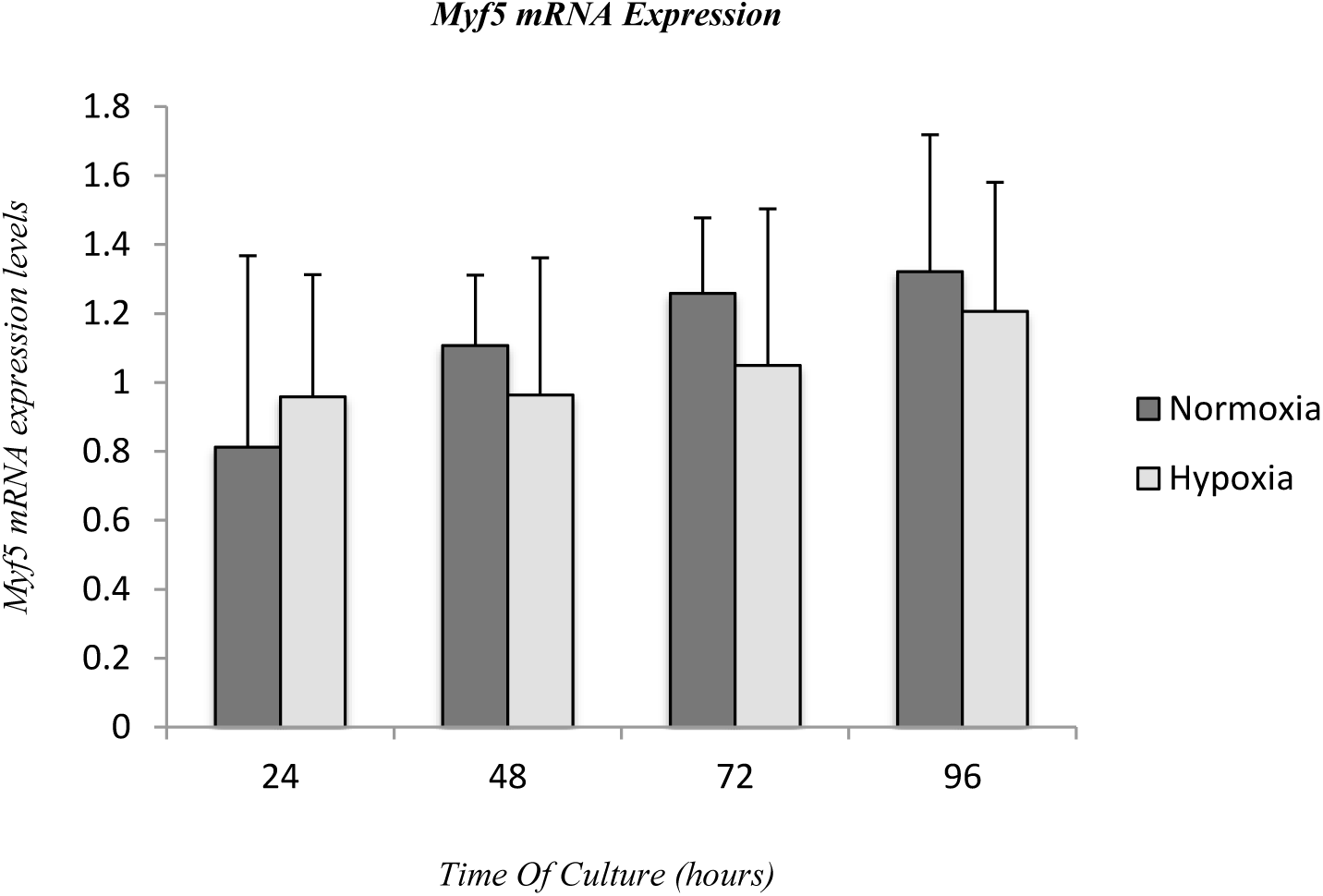
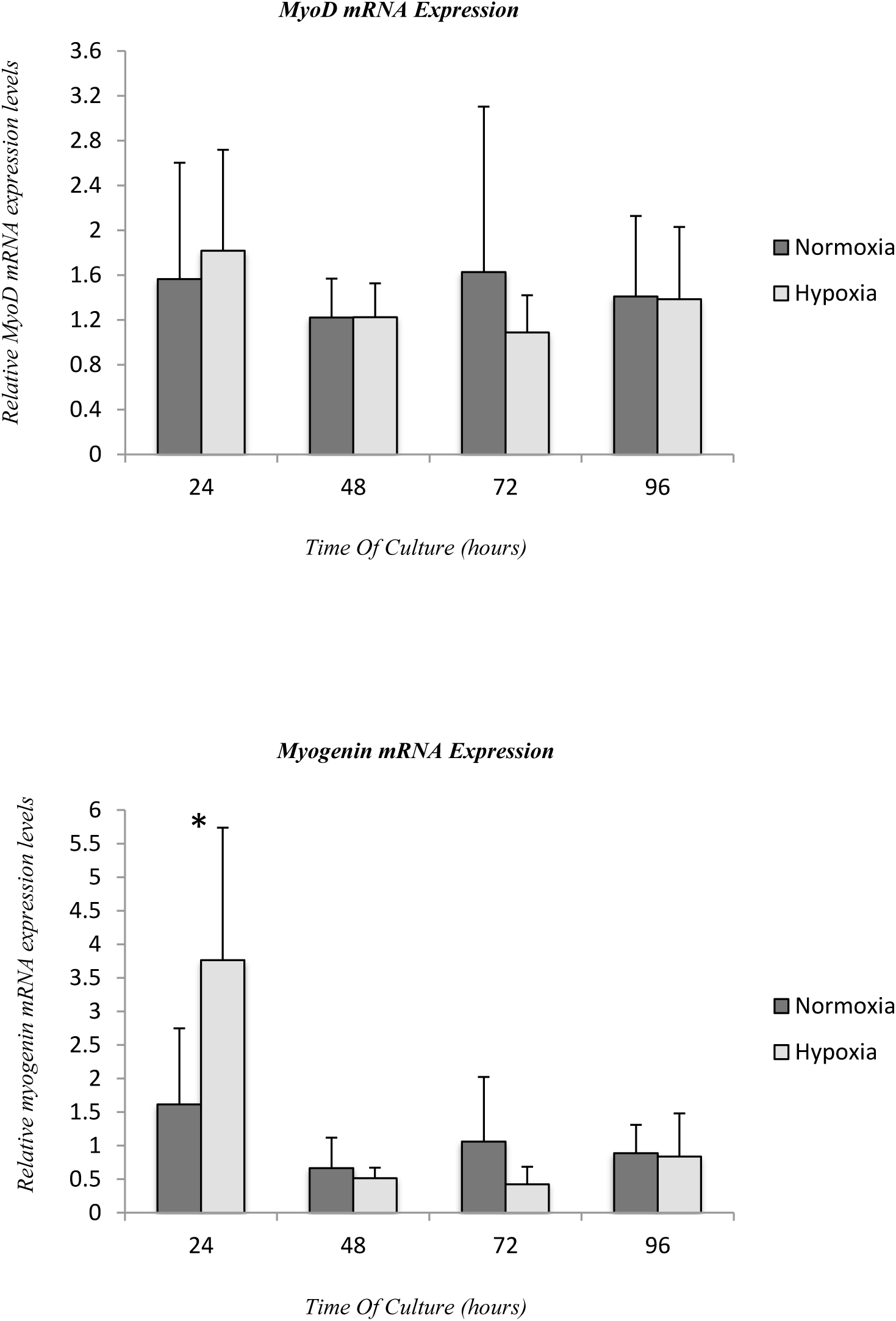
Relative Myf5, MyoD and Myogenin mRNA expression levels from 2D C2C12 cultures, taken every 24 hours over a 96 hour period in vitro, as determined by qPCR. CT values were normalised to an internal housekeeping gene (RPII) and expressed relative to levels recorded following 24 hours of culture in normoxic conditions n=6. Significant difference between timepoints at *P<0.05. Values expressed as mean ± standard deviation.. No significant differences were observed between Normoxic and Hypoxic exposure at any time point with all three genes. (P >0.05)

**Figure. 9.**
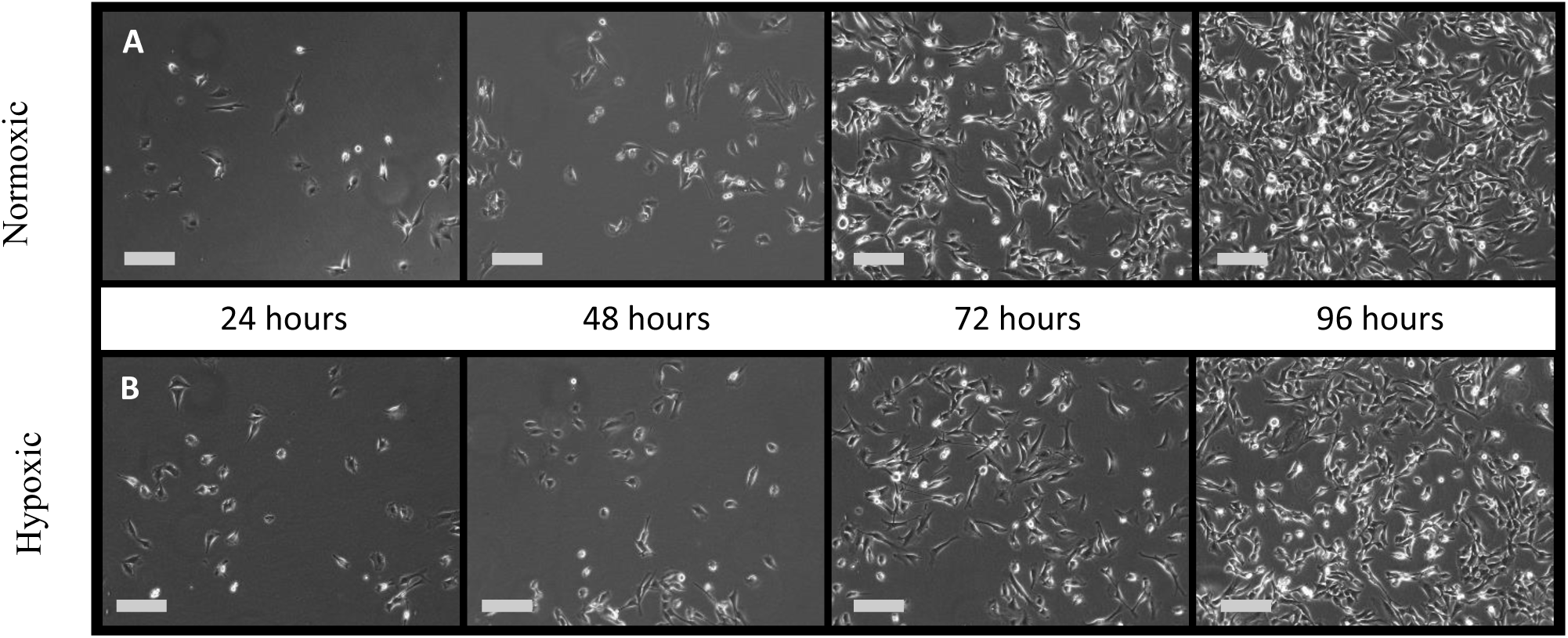
Murine C2C12’s *in vitro*. C2C12 cells were cultured in GM for a total of four days. Cells were fixed every 24 hours. (A) Cells incubated under normoxic conditions (20% O_2_) (B) Cells incubated under normoxic conditions (20% O_2_) and exposed to acute hypoxic exposure (5% O_2_) during the final 24 hours of culturing. Images selected randomly from a selection of ten. Scale bar = 20µm.

## DISCUSSION

The aim of this study was to establish whether acute low oxygen exposure influences the proliferation rate, viability and MRF expression of C2C12 cells *in vitro*. This study has shown that C2C12 myoblasts are unaffected following acute exposure to a low oxygen environment during the proliferative phase of myogenesis. The hypoxic exposure which involved exposure of 5% O_2_ during the final 24hours of culturing upto and including 96hours of total culture time, elicited no significant change in proliferative rate, cell viability or mRNA expression of *Myf5, MyoD* and *Myogenin* compared to cells grown in atmospheric conditions. Further analysis of these images also indicated no myotube formation had occurred following both the normoxic and hypoxic treatments.

It is generally accepted that most stem cells that are cultivated in lower, physiologic oxygen environments, proliferate more than in 21% O_2_ environments (Ramirez, 2011, Csete, 2005; Li *et al*, 2007). This improvement in proliferative capacity in low oxygen has been found in a variety of stem cells including, murine CNSderived multipotent stem cells (Studer *et al*, 2000), marrow-derived mesenchymal stem cells (Lennon *et al*, 2001), and adult murine skeletal muscle satellite cells (Csete *et al*, 2001; Chakravarthy *et al*, 2001). Regarding the cultivation of skeletal muscle satellite cells, the increase in proliferative capacity has been found when satellite cells were cultured between 2% and 10% pO_2_. The enhanced proliferation of stem cells in lower oxygen may in part reflect the typical relatively low oxygen microenvironment that the satellite cell are found *in vivo* which is usually around 2-10% pO_2_ (Greenbaum *et al*, 1997; Richardson, 1998). Although the C2C12 cells used in this study are a mouse myoblast cell line typically cultivated in atmospheric conditions. Therefore 21% O_2_ could be considered normoxic whilst 5% O_2_ would be deemed extremely hypoxic for C2C12 cells. Generally, the increase in proliferation rate elicited is believed to be down to the effect the change in O_2_ has on the signalling pathways that regulate stem cell proliferation and MRF expression. Specifically chronic low oxygen exposure (2-10% pO_2_) has been found to result in a significant up-regulation of *MyoD, Myf5* and *Myogenin* in a number of skeletal muscle cell lines (Yun et al, 2005, Kook et al, 2008, Csete et al 2001).

This experiment found no change in proliferative capacity or MRF expression when cells were cultured in a low oxygen environment, although this is the first experiment of its kind to look at the effects of acute hypoxic exposure (24hours) at different time points during the proliferative phase of myogenesis, rather than chronic exposure. The design study aimed to establish whether acute exposure is effective at eliciting a change in proliferative capacity or MRF expression and if so then at which stage of the proliferative phase of myogenesis should it be used. Sato *et al* (2011) looked at the effect of culturing C2C12’s continuously for 122hours at 2%, 5% and 10% oxygen levels. Interestingly they found that growth rate of the cells in the hypoxic environments was the same as cells under normal conditions for upto 2 days after plating before growth rate increased (Sato *et al*, 2011). Lui *et al* (2012) found that the initial 48hours of culturing primary murine myoblasts in 1% pO_2_ affected myoblast fate decision by promoting self-renewal and inhibiting differentiation, but overall cell proliferation was not affected. However, earlier research by Csete *et al* (2001) did still find a significantly improved rate of proliferation of satellite cells found on cultured single muscle fibres following as little as 24hours of hypoxic exposure at 6% O_2_. Nevertheless, the results of this study with that of recent literature, do tend to indicate 24hours of exposure to a low, physiological oxygen environment may not be an adequate exposure time to induce an effect on cell proliferation rate. It is still unclear whether the hypoxic condition used in the present study reflects the drop in intramyocellular oxygen pressure induced by exercise, muscular damage or hypoxic exposure (Sato *et al*, 2011; Hoppeler *et al*, 2003), however when using manipulation of O_2_ concentrations in improving muscle self renewal for damage repair or treating regenerative disease, it may be possible that exposure times must be longer than 24hours to be effective.

Concerning the effects acute low oxygen exposure has on MRF expression, this study found no difference over time with regards to *Myf5, MyoD* and *Myogenin* expression. Aswell as chronic low oxygen exposure being known to increase expression of all these transcription factors, short term exposure has been shown to upregulate the myogenic regulatory factors *Myf5* and *MyoD* when at 6% O_2_ (Csete *et al*, 2001). When culturing cells in more extreme hypoxic conditions (2 -0.1% pO_2_), there appears to be a down regulation of *Myf5* and *MyoD* expression (Sato *et al*, 2011; Koning *et al*, 2011, Lui *et al* 2012). As no research has yet looked at the effects acute exposure at 5% O_2_ has on these specific MRF’s, it may be possible that 5% O_2_ is the cut off point between which hypoxia either starts having an up-regulatory or down-regulatory effect on *Myf5* and *MyoD* and expression. Concerning *myogenin*, its expression has been shown not to differ significantly by cells under normoxic or hypoxic environment for upto 96hours after induction (Sato *et al*, 2011) supporting our findings and indicating as to why we did not see a significant change in *myogenin* after such a short hypoxic exposure. However *myogenin*, a transcription factor that stimulates myoblasts to undergo terminal differentiation and fusion into myofibers, typically does not undergo up-regulation until both *Myf5* and *MyoD* are coexpressed, usually around 96 hours after plating of C2C12’s (Rudnicki *et al*, 2008; Sato *et al*, 2011, Yablonka-Reuveni and Rivera 1994). The findings of this study found *myogenin* expression was significantly higher following 24hours of culturing in both normoxic and hypoxic conditions, before a rapid down-regulation and no change between 48-96hours of culturing (Figure.8). Explanations for this occurrence are challenging.

Regarding cell viability, this study found that 24hours of acute physiologically hypoxic exposure did not influence cell viability during the proliferative phase of C2C12’s culturing i*n vitro*. This is in contrast to the research looking at the effects of chronic physiological hypoxic exposure that has found various stem cells undergo significantly less apoptosis of cultures in low oxygen than in the traditional 21% O_2_ conditions (Erkkila *et al*, 1999, Csete *et al*, 2001). Furthermore the usual reduction in apoptosis has been found to lead to the increased yield of cultured stem cells during the proliferative phase of cells cultured within low oxygen conditions (Studer *et al*, 2000). Specifically looking at murine muscle satellite stem cells, C2C12’s undergo significantly less apoptosis in 6% O_2_ than in 20% O_2_ following 3 days of exposure (Csete et al, 2001). The possible mechanisms suggested for the anti-apoptotic effect of culturing in low oxygen conditions, is that fewer reactive oxygen species are generated in cells cultured in low oxygen conditions, compared to higher oxygen conditions. Specifically primary murine myoblasts cultured in 6% pO_2_ generated around 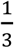 fewer reactive oxygen species compared to cells cultured in 20% oxygen (Csete, 2005). On the contrary Ramires *et al* (2011) found following three days of culturing in 5% pO_2_ there is a significant rise in oxidative stress leading to increased apoptosis. It must be reiterated that these studies looked at the effect of chronic hypoxic exposure, with at least 48hours of exposure. This study used similar oxygen concentrations during the hypoxic exposure, but the exposure time during this study was only 24hours and deemed acute. Therefore the reason that no difference in cell viability was found, leads to the possible suggestion that acute exposure does not allow for an adequate change in reactive oxygen species concentrations and therefore an effect on cell viability. Research that has looked at the apoptosis of neural stem cells in hypoxic conditions, did not find a significant difference until 72hours of exposure in 1%, 2.5% and 5% pO2 (Santilli et al, 2010).

Analysing the techniques used to assess cell viability, it appears the methods used could have been performed more effectively to give a more accurate analysis of cell viability. A heamoctyometer using the Tyrpan blue exclusion method was used for cell counts, yet this would only have been looking at the cells still adhered to the bottom of the six well plate. Although a PBS wash was not performed prior to cell counts to prevent the removal of non-viable cells, the growth medium was removed and any non-adhered cells, most likely the non-viable cells, would not have been accounted for. Therefore an accurate count of the non-viable cells would not have been performed. Future research should aim to include non-adhered cells within the growth medium in the cell counts by centrifuging the removed medium and re-suspending these cells prior to total cell counts being performed, leading to a more accurate account of cell viability and proliferation rates. Independent of this observation, when analysing the cell viability of adhered cells, it can be suggested that the acute hypoxic exposure did not influence the percentage of viable cells as well as the proliferation rate.

It should be noted that the physiologic hypoxic environment in which the cells were cultured (5%pO_2_) was regulated using the Sanyo O_2_/CO_2_ Incubator. Although oxygen sensors embedded within the incubator would ensure the atmospheric environment inside were kept at a constant, it is also important to ensure the oxygen tension seen at the level of the tissue culture monolayer is also expressive of a physiologic hypoxic environment. The oxygen tension seen at a cellular level can vary considerably to the level of oxygen that the incubators deliver, and a number of factors can influence this variation, including the depth of medium, the density of cells, and cellular respiration (Csete, 2005; Muschler, Nakamoto & Griffith, 2004). Within this study to monitor the oxygen tension seen by the cells, a hole was drilled into the top of a single well of the plates and a dip-type oxygen microelectrode was placed within the growth medium at the cellular level. This approach to incorporate multiple oxygen sensors embedded at various levels to monitor oxygen tension is a novel approach that is yet to be seen within studies that culture C2C12 in hypoxic environments. The results indicated that the oxygen tension seen at the cell level was also 5% pO_2_ at the end of the twenty-four hour exposure time in the hypoxic incubator and it can be confirmed the cells were exposed to a physiologically hypoxic environment..

Conversely, it should be noted that due to the fact oxygen tension was only measured at the end of the 24hours of hypoxic exposure within the incubator, a full expression of the change in oxygen tension throughout the twenty-four hour exposure time is not given. Therefore, it cannot be known how long it took the medium to drop to the 5% pO_2_ or whether the exposure time differed as the cells became more confluent. Future research, should try to monitor cell level oxygen tension throughout the hypoxic exposure time to establish at which point a 5% pO_2_ is attained, and also the probe should be placed within the cells cultured in the normoxic environment to ensure those cells are exposed to a normoxic environment. A further limitation is that although the incubator allowed control of oxygen levels during cultivation, it is impossible to prevent reperfusion of atmospheric conditions when the cells are removed from the incubators before being fixed (Csete, 2005). In order to ensure the cells are exposed to relatively constant levels of oxygen, it would be preferential for cells to be cultured and manipulated in a closed workstation that allows for oxygen and CO_2_ control.

Although a considerable amount of work has looked at the effects exercise and altitude have on oxygen concentrations within the body (Howald & Hoppeler *et al*, 2003; Hoppeler *et al*, 2008), it is still unclear whether the hypoxic condition used in the present study reflect the drop in intramyocellular oxygen pressure induced by exercise, altitude or pathological damage (Sato *et al*, 2011; Hoppeler *et al*, 2003). Therefore the results of this study may not yet be related to the effects exercise or acute altitude training have on skeletal muscle proliferation rates within humans. Although should intramyocellular oxygen pressures be able to be manipulated, it may be possible to suggest twenty-four hours of exposure at 5% O_2_ is inadequate in inducing improved muscular regeneration. That being said, *in vitro* it can be concluded that the effect of acute low oxygen exposure lasting 24hours appears to have no effect on the proliferation rate, viability or transcription factor expression of C2C12’s during the proliferative phase of myogenesis. Future research may wish to use more stringent methods with regards to fixing cells within a hypoxic environment and better controls during counts of cell viability. Furthermore, a look at whether lower concentrations of O_2_ on the proliferative capacity of C2C12’s, aswell as various exposure times is needed to develop a greater understanding of the effects acute hypoxia has on the proliferative phase of myogenesis.

